# Symbiont-mediated fly survival is independent of defensive symbiont genotype in the *Drosophila melanogaster-Spiroplasma*-wasp interaction

**DOI:** 10.1101/2020.07.05.154906

**Authors:** Jordan E. Jones, Gregory D. D. Hurst

**Affiliations:** Institute of Infection, Veterinary and Ecological Sciences, University of Liverpool, Liverpool, L69 7ZB, UK

**Keywords:** *Drosophila melanogaster*, symbiont-mediated protection, *Leptopilina*, *Spiroplasma*

## Abstract

When a parasite attacks an insect, the outcome is commonly modulated by the presence of defensive heritable symbionts residing within the insect host. Previous studies noted markedly different strengths of *Spiroplasma*-mediated fly survival following attack by the same strain of wasp. One difference between the two studies was the strain of *Spiroplasma* used. We therefore performed a common garden laboratory experiment to assess whether *Spiroplasma*-mediated protection depends upon the strain of *Spiroplasma*. We perform this analysis using the two strains of male-killing *Spiroplasma* used previously, and examined response to challenge by two strains of *Leptopilina boulardi* and two strains of *Leptopilina heterotoma* wasp. We found no evidence *Spiroplasma* strain affected fly survival following wasp attack. In contrast, analysis of the overall level of protection, including the fecundity of survivors of wasp attack, did indicate the two *Spiroplasma* strains tested varied in protective efficiency against three of the four wasp strains tested. These data highlight the sensitivity of symbiont-mediated protection phenotypes to laboratory conditions, and the importance of common garden comparison. Our results also indicate that *Spiroplasma* strains can vary in protective capacity in *Drosophila*, but these differences may exist in the relative performance of survivors of wasp attack, rather than in survival of attack per se.

## Introduction

The outcome of natural enemy attack has traditionally been considered a function of factors encoded within the genome of the host and infecting parasite. Within this interaction may exist a degree of specificity whereby a subset of parasite genotypes are able to infect a subset of host genotypes and, reciprocally, a subset of host genotypes are able to resist a subset of parasite genotypes (Woolhouse *et al*. 2002; Lambrechts *et al*. 2006). Specificity between host and parasite genotypes can lead to negative-frequency dependent selection between players and can contribute to the maintenance of heritable variation for defence and attack factors within a population (Woolhouse *et al*. 2002; Schmid-Hempel & Ebert 2003).

More recently it has been observed that the outcome of natural enemy attack is not solely determined by host and parasite genotypes, but also by the presence and genotype of defensive heritable microbial symbionts residing within the host (Brownlie & Johnson 2009; Oliver *et al*. 2009; Ballinger & Perlman 2019). Defensive microbial symbionts have been identified in a wide range of organisms. For example, microbial symbionts are known to provide protection against ssRNA viruses (Hedges *et al*. 2008; Teixeira *et al*. 2008), nematodes (Jaenike *et al*. 2010), fungal pathogens (Scarborough *et al*. 2005; Lukasik *et al*. 2013) and parasitic wasps (Oliver *et al*. 2003; Xie *et al*. 2010).

Recently, studies have described how microbial strain identity can complement host and parasite genotype as an additional driver of the outcome of a host – parasite interaction. In aphid systems, this is commonly manifested in symbiont strain x host strain x enemy strain interaction terms (Sandrock *et al*. 2010; Schmid *et al*. 2012; Cayetano & Vorburger 2013, 2015; Parker *et al*. 2017). Beyond the aphid systems, it is known that the strain of infecting *Wolbachia* is an important source of variation in *Wolbachia-mediated* protection against viruses in *Drosophila*, associated with different titre achieved by the strains (Osborne *et al*. 2009; Bian *et al*. 2013; Chrostek *et al*. 2013, 2014; Martinez *et al*. 2017). Similarly, in the bumblebee, *Bombus terrestris*, the defensive gut microbiota type is predominantly responsible for resistant phenotypes against the virulent gut trypanosomatid, *Crithidia bombi* (Koch & Schmid-Hempel 2012).

The heritable endosymbiont *Spiroplasma*, has been shown to protect *Drosophila* from attack by nematodes and parasitoid wasps (Jaenike *et al*. 2010; Xie *et al*. 2010, 2014; Mateos *et al*. 2016). The ability of *Spiroplasma* to protect *Drosophila* is thought to be orchestrated through a combination of RIP toxin activity (secreted by *Spiroplasma*) and exploitative competition between *Spiroplasma* and the infecting parasite for lipid stores (Paredes *et al*. 2016; Ballinger & Perlman 2017). Despite being regarded as an important model system, little is known about the role of host, symbiont and parasite identity in determining the outcome of the interaction. Recent work has revealed that the genotype of attacking parasitoid wasp is important for the degree of protection conferred by *Spiroplasma* (Jones & Hurst 2020). It was observed that *Spiroplasma* (MSRO-Br strain) conferred protection of 40% against the Lh-Fr and Lh-Mad *L. heterotoma* wasp strains, contrasting with 5% protection against the Lh14 strain. The reasons underpinning the variation observed is unknown, but intraspecific differences in the toxicity of wasp venom transferred along with the wasp egg during parasitization may be a factor.

A more general understanding of how symbiont and parasite genotypes are likely to interact is essential for predicting the dynamics of symbionts in natural populations. In this study, we determine whether parasite genotype x symbiont genotype interactions exist within the *Spiroplasma*-*Drosophila melanogaster* system. Most studies concerning *Spiroplasma*-mediated protection have reported the outcome of experiment in which a single symbiont strain defends against a single enemy strain. Analysis across these studies indicates that the strain of *Spiroplasma* may be an important component of *Spiroplasma*-mediated protection. For instance, survival of flies exposed to the Lb17 strain of the specialist parasitoid wasp *L. boulardi* was recorded at 5% in *D. melanogaster* infected with the MSRO-Br strain (Xie *et al*. 2014), and 50% in *D. melanogaster* infected with the MSRO-Ug *Spiroplasma* strain (Paredes *et al*. 2016). One interpretation of these results is that the *Spiroplasma* strains differ in protective capacity in *D. melanogaster*. However, analysis of these two strains within a common experimental design (controlling for potential lab practice, wasp strain and fly strain differences) is required to determine the precise importance of symbiont strain in determining the outcome of the parasite-host interaction.

We here present an analysis of the capacity of MSRO-Br and MSRO-Ug to defend *D. melanogaster* against wasp attack. This analysis is performed for two strains of the specialist parasitoid *L. boulardi*, and two strains of the generalist *L. heterotoma*. We compare survival following wasp attack, mirroring previous studies, and additionally estimate overall protection combining fly survival data with data on the fertility of flies that survived wasp attack to establish a protective index for each wasp strain by *Spiroplasma* strain combination.

## Materials and methods

### Strains and maintenance

Two strains of *Spiroplasma* were used in this study. The first, Red 42, was originally collected in Campinas, São Paulo State, Brazil in 1997 (Montenegro *et al*. 2000) and later transinfected and maintained in the laboratory on a Canton-S background. The second *Spiroplasma* strain was collected from Namulonge, Uganda in 2005 (Pool *et al*. 2006) which was later transferred and maintained in the laboratory on an Oregon-R background. It should be noted that all larvae from the *Spiroplasma* infected treatments are female due to the high efficiency of male-killing. However, there does not appear to be any differences in survival between the sexes against parasitoid wasp attack (Xie *et al*. 2014). All flies were maintained on ASG corn meal agar vials (10 g agarose, 85 g sugar, 60 g maize meal, 40 g autolysed yeast in a total volume of 1 L, to which 25 mL 10% Nipagin was added) at 25 °C on a 12:12 light:dark cycle.

The *L. boulardi* strains used were the NSRef strain, established from an initial female collected in Gotheron, near Valence, France (Varaldi *et al*. 2003), and the Lb17 strain, initially collected in Winters, California in 2002 (Schlenke *et al*. 2007). The *L. heterotoma* strains used were the inbred Lh14 strain also collected in Winters, California in 2002 (Schlenke *et al*. 2007) and the Lh-Mad strain established from a single female collected in Madeira, Portugal in March 2017 (Jones & Hurst 2020). The wasp stocks were all maintained on second instar Oregon-R larvae at 25°C on a 12:12 light:dark cycle. After emergence, wasps were maintained on grape agar vials supplemented with a flug moistened with honey water and allowed to mature and mate for 7 days prior to exposure to *D. melanogaster* L2 larvae.

### Artificial infection of *Spiroplasma*

The *Spiroplasma* strains (MSRO-Br and MSRO-Ug) were artificially transferred into a common host background (Canton-S) to remove any effect of host nuclear background on the level of protection conferred. Canton-S stocks carry the naturally occurring *Wolbachia* strain wMel. *Wolbachia* has been shown to provide a weak positive effect on fly larva-to-adult survival and a negative effect on wasp success in flies attacked against *L. heterotoma* (Lh14 strain) (Xie *et al*. 2014). Artificial infections were carried out as described by Nakayama *et al*. (2015). Hemolymph was extracted from the thorax of *Spiroplasma*-infected *D. melanogaster* and mixed with sterile PBS. Virgin female Canton-S were artificially injected in the abdomen with 0.1-0.2 μl of PBS-hemolymph, using a hydraulic positive-pressure microinjection apparatus (Model IM-6, Narushige Ltd, Tokyo, Japan).

### Confirmation of *Spiroplasma* infection status

Three weeks post injection, the infection status of the artificially infected flies was confirmed via *Spiroplasma*-specific PCR. DNA extraction was carried out using the Wizard^®^ Genomic DNA Purification Kit (Promega). To this end, each injected mother was taken and macerated in 150 μl of Nuclei Lysis Solution and incubated at 65 °C for 30 min. After incubation, 50 μl of Protein Precipitation Solution was added to each sample and then placed on ice for 5 min. Samples were then centrifuged for a further 4 min at 16,000 x g and the supernatant was transferred into a new tube containing 150 μl of isopropanol. Samples were centrifuged for 2 min at 16,000 x g and the supernatant discarded. 150 μl of 70% ethanol was added to each sample and centrifuged for 1 min at 16,000 x g. The supernatant was discarded. Pellets were dried before re-suspending in 25 μl of molecular grade water at 4 °C overnight before use in subsequent PCR assays.

PCR amplifications were conducted using *Spiroplasma* specific primers, SpoulF (5’-GCT TAA CTC CAG TTC GCC-3’) and SpoulR (5’-CCT GTC TCA ATG TTA ACC TC-3’) (Montenegro *et al*. 2005). Each reaction was carried out in 15 μl volume containing 7.5 μl of GoTaq^®^ Hot Start Green Master Mix (Promega), 0.5 μl each of the forward and reverse primer, 5.5 μl of Molecular Grade Water and 1 μl of DNA. All reactions were conducted alongside the positive and negative controls. This included a PCR negative control containing the PCR reaction mixture only (excluding DNA template). The PCR thermal program consisted of an initial denature of 5 min at 95 °C, followed by 35 cycles of 15 s at 94 °C, 1 min at 55 °C and 40 s at 72 °C. The PCR products were electrophoresed in a 1.5% agarose gel at 155 V for 15 min and the products were visualised. Offspring sex ratio of infected mothers were also checked to determine *Spiroplasma* efficiency. Only mothers which were infected with *Spiroplasma* and produced all female broods were used to create new lines.

To confirm the *Spiroplasma* strain status of each artificially injected line of *Drosophila melanogaster*, sequencing was performed on 5 individual flies from each strain. To this end, the DNA of 5 flies from each *Spiroplasma* strain line were extracted using the Wizard^®^ Genomic DNA Purification Kit following the methodology from above. PCR amplifications were conducted using *Spiroplasma* specific primers, Spiro_MSRO_diff_F (5’-TAC GAC CAA TGG CTT GTC CC-3’ and Spiro_MSRO_diff_R (5’-CTG GCA TTG CTT TTT CCC CA-3’). The PCR thermal program consisted of an initial denature of 2 min at 94 °C, followed by 35 cycles of 15 s at 94 °C, 1 min at 56 °C and 40 s at 72 °C. To prepare the PCR reaction for sequencing, PCR products underwent an ExoSAP digest clean up to remove excess primers. To this end, 5 μl of PCR product was added to a mixture containing 0.2 μl Shrimp alkaline phosphate, 0.05 μl of Exonuclease I, 0.7 μl 10X RX Buffer and 1.05 μl of molecular grade water. Samples were then incubated for 45 min at 37 °C followed by 15 min at 80 °C and sent for Sanger sequencing. The *Spiroplasma* strain status of the MSRO-Br and MSRO-Ug line were confirmed by the presence of a Guanine and Thymine respectively in position 414193, coding for a type III pantothenate kinase. The expected amplicon size is 509bp. Transinfected fly lines were passaged for >10 generations before experiments were conducted.

### Wasp attack assay

To ensure efficient vertical transmission of *Spiroplasma*, infected females were aged to at least ten days prior to egg laying. Flies were allowed to mate in cages and lay eggs on a grape Petri dish painted with live yeast for 24 h. Grape Petri dishes were incubated for a further 24 h to allow larvae to hatch. First instar larvae were picked from the grape plate into the experimental vials at 30 larvae per vial. A fully factorial design was created for each of the four wasp strains described which included *Spiroplasma* strain (MSRO-Br, MSRO-Ug and uninfected control) and wasp (presence or absence). Five experienced, mated female wasps were transferred into the wasp treatment vials. Adult wasps were allowed to parasitise for 2 days before being removed. All vials were maintained at 25 °C on a 12:12 light:dark cycle. For each vial, the number of puparia, emerging flies and emerging wasps were recorded. Experiments using *L. boulardi* and *L. heterotoma* were conducted in separate blocks, one week apart.

### Measuring female fecundity

*Spiroplasma* infected flies that survive wasp attack generally have a lower fecundity than *Spiroplasma* infected flies which were not exposed to wasps (Xie *et al*. 2011; Jones & Hurst, 2020). To determine whether the wasp attacked survivors were differentially impacted by *Spiroplasma* strain the average daily emerged offspring of *Spiroplasma* infected survivors (“Exposed”) and *Spiroplasma* infected flies which did not undergo wasp attack (“Unexposed”) was measured for the MSRO-Br and MSRO-Ug strains. The *Spiroplasma* uninfected wasp attacked group was not included due to the extremely low number of flies which emerged, which were also likely to have avoided wasp attack all together. After emergence, flies from the wasp attack assay were stored in vials containing sugar yeast medium (20 g agarose, 100 g sugar, 100 g autolysed yeast in a total volume of 1 L, to which 30 mL 10% Nipagin w/v propionic acid was added) at mixed ages. A week after emergence commenced, a subset of flies from each of the *Spiroplasma* treatments were placed into an ASG vial with two Canton-S males with a single yeast ball and allowed to mate. Approximately 25 replicates per treatment were created. Flies were transferred onto fresh ASG vials each day for five days. Flies were given two weeks to emerge to ensure every fly had emerged before counting. Female fecundity was measured as the average number of offspring produced over four days (day 2-5).

### Statistical analysis

All statistical analyses were performed using the statistical software R, version 3.5.0 (R Core Team 2018). Fly and wasp survival data were analysed by fitting a generalized linear model with binomial distributions. In all cases, a fully saturated model including all factors and their interaction was reduced to a minimum adequate model through step-wise simplification. Non-significant factors are reported as the output of the model comparisons. The effect of significant independent variables are reported from the analysis of the minimum adequate model using the ‘car’ package.

To produce a composite measure of protection, a Protective Index (PI) was calculated by comparing the survival and fecundity of *Spiroplasma*-infected flies in the presence/absence of a given strain of wasp. The PI was calculated as the ratio of p(survival) x p(fertile) x fecundity of fertile individuals for attacked vs unattacked *Spiroplasma*-infected flies and reflects the benefit of *Spiroplasma* in the face of wasp attack. Credible intervals for PI were calculated through simulation. By assuming prior probability distributions for each parameter (Survival probability = beta distribution; Fertility probability = beta distribution; Fecundity = normal distribution), the ‘rbeta’ and ‘rnorm’ functions were used to calculate 95% credible intervals for PI. The simulation data was also used to establish the posterior probability of PI differing between attacking wasp strains.

## Results

### Fly survival and wasp success

#### Leptopilina boulardi experiment

In the absence of *L. boulardi* wasps, *Spiroplasma* strain had a significant effect on fly larva-to-adult *D. melanogaster* survival (χ^2^ = 7.74, d.f. = 1, *P* = 0.005). The mean survival of MSRO-Br infected and MSRO-Ug infected *D. melanogaster* was 72.2% and 83%, respectively (Fig. 1A). In the presence of *L. boulardi* wasps, there was no significant effect of wasp strain (χ^2^ = 0.281, d.f. = 1, *P* = 0.596), *Spiroplasma* strain (χ^2^ = 0.0008, d.f. = 1, *P* = 0.977), nor a significant interaction between wasp strain and *Spiroplasma* strain on larva-to-adult survival of *D. melanogaster* (χ^2^ = 0.284, d.f. = 1, *P* = 0.594) (Fig. 1A). There was no significant effect of wasp strain on wasp success (χ^2^ = 0.121, d.f. = 1, *P* = 0.728) (Fig. 1A), and wasps were observed only in the absence of *Spiroplasma*.

**Figure 1:**
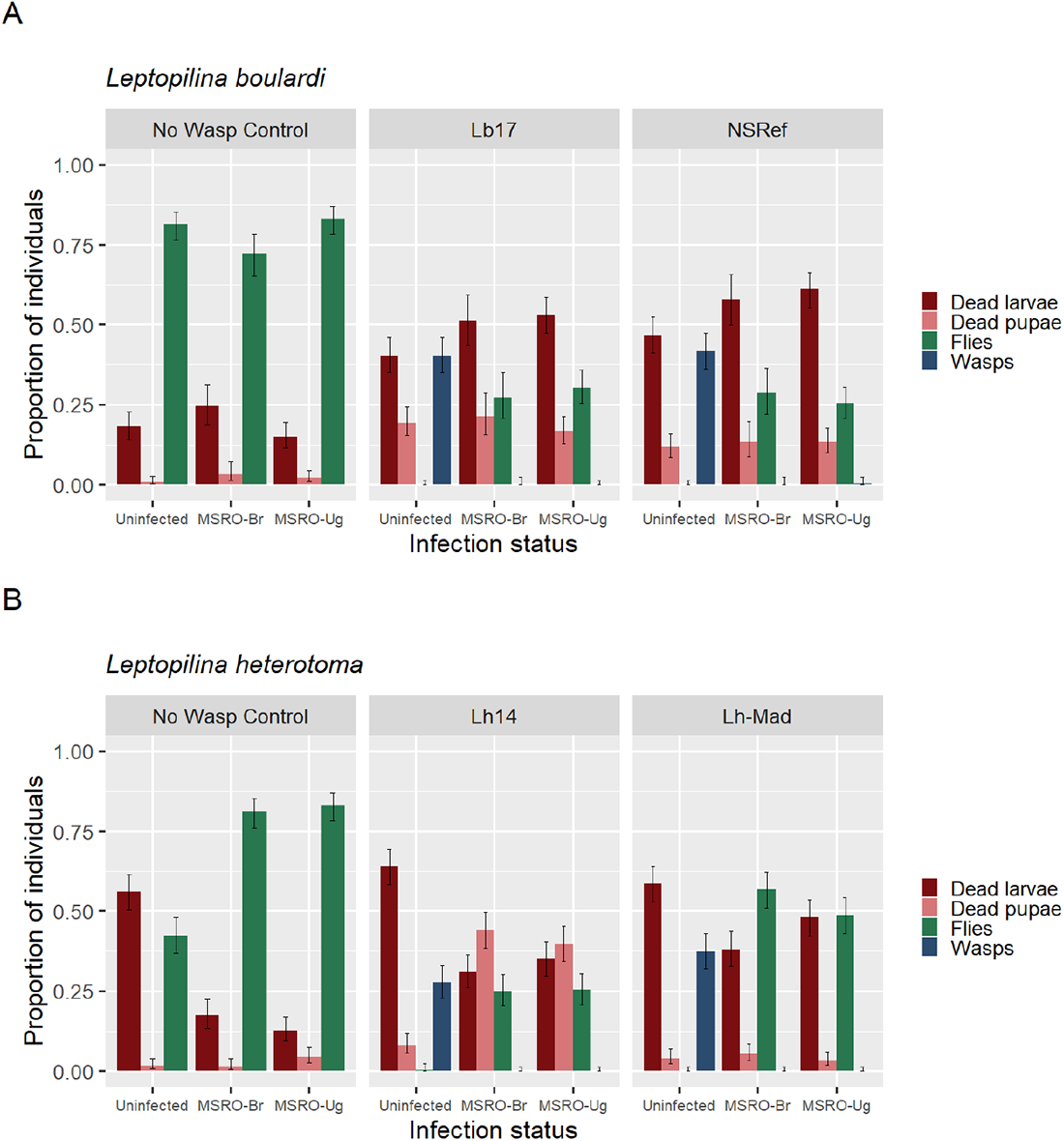
Proportion of dead larvae (red), dead pupae (pink), emerging flies (green) and emerging wasps (blue) for *Spiroplasma*-infected (MSRO-Br and MSRO-Ug strains) and uninfected *Drosophila melanogaster* attacked by A) *L. boulardi* (Lb17 and NSRef strains) and B) *L. heterotoma* (Lh14 strain and Lh-Mad strains). Error bars represent 95% binomial confidence intervals.

#### Leptopilina heterotoma experiment

In the absence of *L. heterotoma* wasps, there was no significant effect of *Spiroplasma* strain on fly larva-to-adult survival (χ^2^ = 0.345, d.f. = 1, *P* = 0.557). The mean survival of uninfected, MSRO-Br infected and MSRO-Ug infected *D. melanogaster* was 81.1% and 83%, respectively (Fig. 1B). In the presence of *L. heterotoma* wasps, there was a significant effect of wasp strain on fly larva-to-adult survival of *D. melanogaster* (χ^2^ = 34.21, d.f. = 1, *P* < 0.001). Fly larva-to-adult survival of *Spiroplasma*-infected *D. melanogaster* attacked by the Lh-Mad strain of *L. heterotoma* was approximately double that observed for flies attacked by the Lh14 strain of *L. heterotoma* (Fig. 1B). Similar to the *L. boulardi* experiment, there was no significant effect of *Spiroplasma* strain (χ^2^ = 0.740, d.f. = 1, *P* = 0.390), nor a significant interaction between wasp strain and *Spiroplasma* strain (χ^2^ = 0.674, d.f. = 1, *P* = 0.412) on larva-to-adult survival of *D. melanogaster* (Fig. 1B). There was a significant effect of wasp strain on wasp success (χ^2^ = 4.805, d.f. = 1, *P* = 0.028) (Fig. 1B). The average wasp success of the Lh14 and Lh-Mad wasp strains were 27.7% and 37.3% respectively. Wasps only emerged in the absence of *Spiroplasma*, with both symbiont strains preventing development of both wasp strains.

### Overall protection index

Despite finding no difference between the survival of flies infected with MSRO-Br and MSRO-Ug against each of the four wasp strains tested, previous work has shown that it is also important to consider, in combination with survival, the fertility of wasp-attacked flies compared to non-attacked controls to produce a complete model of protection (Xie et al., 2011; Jones & Hurst 2020). Taking into account the survival, proportion of adults fertile, and the fecundity of wasp attack survivors, compared to unexposed *Spiroplasma*-infected controls, a protection index (PI) was calculated as the product of fly survival x p(fertile) x fecundity of exposed vs unexposed *Spiroplasma*-infected flies (Table 1). This metric assumes complete mortality from wasps in the absence of *Spiroplasma*, which is approximately true as <1% of individuals tested survived wasp attack. Against the Lb17, NSRef and Lh-Mad strains of wasp, the posterior probability that the protection index for MSRO-Br is greater than the protection index for MSRO-Ug is >0.97 (Table 2). However, against the Lh14 strain of wasp, the posterior probability that the protection index for MSRO-Br is greater than the protection index for MSRO-Ug is 0.44 (Table 2).

**Table 1:** The overall protection conferred by MSRO-Br and MSRO-Ug *Spiroplasma* strains against a) *Leptopilina boulardi* (Lb17 and NSRef strains) and b) *Leptopilina heterotoma* (Lh14 and Lh-Mad strains) in *Drosophila melanogaster*. Exposed S- = wasp attacked *Spiroplasma*-uninfected flies; Exposed S+ wasp attacked *Spiroplasma*-infected flies; Unexposed S+ *Spiroplasma*-infected flies not attacked. Protective Index is calculated as [p(survival) x p(fertile) x fecundity of fertile individuals] of exposed vs unexposed individuals with credible intervals calculated as given in methods.

| *Spiroplasma* strain | Treatment | Fly Survival (binomial 95% CI intervals (lower, upper)) | Proportion fertile (binomial 95% CI intervals (lower, upper)) | Fecundity measure ± SE | Estimated protective index (95% Credible interval (lower, upper)) |
| --- | --- | --- | --- | --- | --- |
| MSRO-Br | Lb17 exposed S+ | 0.27 (0.20, 0.35) | 0.96 (0.75, 0.99) | 15.8 ± 1.31 | 0.37 (0.25, 0.55) |
| | NSRef exposed S+ | 0.29 (0.22, 0.36) | 1.00 (0.86, 1.00) | 15.6 ± 1.45 | 0.40 (0.27, 0.59) |
| | Unexposed control S+ | 0.72 (0.65, 0.78) | 0.92 (0.72, 0.98) | 16.9 ± 1.47 | |
| MSRO-Ug | Lb17 exposed S+ | 0.30 (0.25, 0.36) | 1.0 (0.85, 1.0) | 11.0 ± 1.77 | 0.30 (0.14, 0.32) |
| | NSRef exposed S+ | 0.25 (0.21, 0.30) | 0.88 (0.68, 0.96) | 14.3 ± 1.58 | 0.20 (0.14, 0.30) |
| | Unexposed control S+ | 0.83 (0.78, 0.87) | 0.96 (0.776, 0.99) | 19.4 ± 1.49 | |

b) *Leptopilina heterotoma*
| *Spiroplasma* strain | Treatment | Fly Survival (binomial 95% CI intervals (lower, upper)) | Proportion fertile (binomial 95% CI intervals (lower, upper)) | Fecundity measure ± SE | Estimated protective index (95% Credible interval (lower, upper)) |
| --- | --- | --- | --- | --- | --- |
| MSRO-Br | Lh14 exposed S+ | 0.25 (0.20, 0.30) | 0.79 (0.59, 0.91) | 13.2 ± 1.75 | 0.24 (0.15, 0.39) |
| | Lh-Mad exposed S+ | 0.57 (0.51, 0.62) | 0.91 (0.71, 0.98) | 14.0 ± 1.58 | 0.68 (0.54, 1.16) |
| | Unexposed control S+ | 0.81 (0.76, 0.85) | 0.92 (0.71, 0.98) | 14.3 ± 1.72 | |
| MSRO-Ug | Lh14 exposed S+ | 0.25 (0.20, 0.31) | 1.0 (0.82, 1) | 12.8 ± 1.14 | 0.25 (0.18, 0.36) |
| | Lh-Mad exposed S+ | 0.49 (0.43, 0.54) | 0.91 (0.70, 0.98) | 11.0 ± 1.34 | 0.39 (0.26, 0.56) |
| | Unexposed control S+ | 0.83 (0.78, 0.87) | 1.0 (0.82, 1) | 15.3 ± 1.58 | |

**Table 2:** The posterior probability that the estimated protective index for MSRO-Br is greater than the MSRO-Ug for each wasp strain tested.

| Wasp strain | Estimated protective index (95% Credible interval<br>(lower, upper)) |  | Posterior probability (EPI<br>MSRO-Br > EPI MSRO-Ug) |
| --- | --- | --- | --- |
|  | MSRO-Br | MSRO-Ug |  |
| *Leptopilina boulardi* |  |  |  |
| Lb17 | 0.37 (0.25, 0.55) | 0.30 (0.14, 0.32) | 0.97 |
| NSRef | 0.40 (0.27, 0.59) | 0.20 (0.14, 0.30) | 0.99 |
| *Leptopilina heterotoma* |  |  |  |
| Lh14 | 0.24 (0.15, 0.39) | 0.25 (0.18, 0.36) | 0.44 |
| Lh-Mad | 0.68 (0.54, 1.16) | 0.39 (0.26, 0.56) | 0.99 |

## Discussion

Defensive symbionts can contribute to the outcome of a host-parasite interaction. Previous studies in aphids have shown that the strain of symbiont is an important determinant of symbiont-mediated protection across multiple model systems (Schmid *et al*. 2012; Cayetano & Vorburger 2013, 2015; Parker *et al*. 2017). However, whether strains of the *Drosophila* defensive symbiont, *Spiroplasma poulsonii*, vary in their capacity for protection is unknown. The contrasting levels of fly survival observed between two previous studies on the *Drosophila-Spiroplasma-L. boulardi* interaction suggested that the strain of *Spiroplasma* may be an important determinant of protection capacity in *Drosophila* (Xie *et al*. 2014; Paredes *et al*. 2016). We therefore performed an experiment to determine whether the strength of *Spiroplasma*-mediated protection depended on the strain of infecting *Spiroplasma* using two known strains of MSRO *Spiroplasma* (MSRO-Br and MSRO-Ug). We found no evidence that the strength of *Spiroplasma*-mediated fly survival differed between the MSRO-Br and MSRO-Ug strains against any of the four *Leptopilina* wasp strains tested. However, the overall protective index, including the fecundity of survivors of wasp attack, did vary between the two *Spiroplasma* strains for three of the attacking wasp strains.

The strain of *Spiroplasma* did not alter the strength of *Spiroplasma*-mediated fly survival in *D. melanogaster* in our experiment. This result raises the question as to why fly survival following attack differed between the two previous independent studies. Fly survival against the parasitoid wasp, *L. boulardi* (strain Lb17) was observed to vary from 5% with MSRO-Br (Xie *et al*. 2014), to 50% with MSRO-Ug (Paredes *et al*. 2016). Comparisons across studies indicate that the strength of symbiont-mediated fly survival appear to be highly variable across laboratory studies. In this study, we found survival of 30% against the *L. boulardi* (Lb17 strain), yet Paredes *et al*. (2016) found survival of 50% against the same wasp strain despite using the same fly strain. Similarly, we found survival of 25% against the Lh14 strain of *L. heterotoma*, despite survival of <8% observed in previous studies (Xie *et al*. 2014, Jones & Hurst 2020).

The variability in *Spiroplasma*-mediated survival observed across studies may be the result of variability in wasp success. Whilst wasp attack rate was very high in all cases (with very low fly survival in uninfected controls), wasp success was highly variable across the studies and correlated to some extent with fly survival. Specifically, against the Lb17 wasp strain, Xie *et al*. (2014) found high wasp success of ~70% and low fly survival of ~5%. In contrast, this study observed reduced wasp success of ~40% and increased fly survival of ~30%. Thus, the variability in *Spiroplasma*-mediated fly survival across studies could be associated with condition of the attacking wasps. Associated with this, it is notable that larval-to-pupa survival following attack is lower in our studies than previously observed, and this may potentially explain differences in wasp survival. These studies may highlight the sensitivity of symbiont-mediated protection to husbandry conditions of both fly and wasp.

From several studies, it has been demonstrated that symbiont-mediated survival against natural enemies can be highly sensitive to particular environmental conditions. Temperature is one environmental factor known to impact the strength of symbiont-mediated protection (Corbin *et al*. 2017). For example, in the pea aphid, higher temperatures can negatively impact *H. defensa*-mediated survival against *Aphidius ervi* (Doremus *et al*. 2018). Similarly, heat shock also negatively impacts X-type-mediated survival against *A. ervi* wasps in the pea aphid (Heyworth & Ferrari 2016). Another possibility, raised by studies of the strength of CI and male-killing exhibited by *Wolbachia*, is that protection strength is influenced by parental *Spiroplasma* titre (Dyer *et al*. 2005; Layton *et al*. 2019). It is notable that both thermal environment and age at reproduction are known to affect *S. poulsonii* titre and male-killing strength in *D. melanogaster* (Anbutsu & Fukatsu 2003; Montenegro & Klaczko 2004; Anbutsu *et al*. 2008). Finally, wasp husbandry and attack protocols may vary. Wasp attack success is thought to be higher when wasps are previously conditioning before assays and may also be impacted by the arena in which attack occurs. Wasps attack fly larvae at the surface of the food, and the surface area available for attack, and indeed the medium in which the larvae are feeding, may impact success. The variable strength of protection afforded by symbionts across laboratories may be due to unmeasured differences in stock maintenance/ambient environmental conditions and reinforce the need for common-laboratory experiments when comparing outcomes.

Our experiment nevertheless did indicate differences in protection associated with *Spiroplasma* strain, but these were reflected in the overall phenotype, including the survival and fecundity of wasp-attack survivors. Surviving flies infected with the MSRO-Br strain of *Spiroplasma* had an overall higher protective index against the NSRef, Lb17 and Lh-Mad strains of wasp compared to flies infected with the MSRO-Ug strain. The reasons as to why fly survivors infected with MSRO-Ug had a lower protective index compared to MSRO-Br survivors remains unclear. One possible factor which cannot be ruled out from this study is the effect of *Wolbachia*. Although from the results it does not appear that *Wolbachia* is having an effect on fly survival, it may be possible that the presence of *Wolbachia* is differentially impacting the fertility of wasp-attacked survivors among the MSRO-Br and MSRO-Ug strains tested. Another factor which is difficult to determine is the possibility that a proportion of flies in the *Spiroplasma* treatments were not attacked. Although fly emergence from the *Spiroplasma* negative controls suggests that all larvae were successfully parasitized, this does not exclude the possibility that not all larvae were parasitized in the *Spiroplasma* positive treatments, although past work found no evidence for discrimination by wasps (Xie *et al*. 2010, Jones & Hurst, 2020). However, the result that there was no difference in the overall protection between wasp-attacked survivors infected with MSRO-Br and MSRO-Ug against the Lh14 strain of wasp indicates that the reasons for this difference may be a consequence of wasp strain.

This study clearly demonstrates two important features of protection. First, there is a need for common-laboratory experiments to compare levels of protection, as this phenotype has both genetic and environmental drivers. Second, there is a clear distinction between symbiont-mediated survival and symbiont-mediated protection within defensive symbiont studies. Symbiont-mediated protection is often measured as the relative survival of an infected-individual compared to an uninfected individual when faced with natural enemy attack. However, symbiont-mediated protection is not only the ability of an infected-host to survive, but also the relative capacity it has to reproduce compared to un-attacked comparators. Despite finding no evidence that fly survival differed between the two strains of *Spiroplasma* against all four wasp strains tested, differences between *Spiroplasma* strains were observed on the overall strength of symbiont-mediated protection. Assessment of the relative survival and reproductive ability of un-attacked vs. attacked survivors is essential for revealing the true protective capacity of a defensive symbiont.

## Acknowledgments

We would like to thank Dr Florent Masson for providing the MSRO-Ug *D. melanogaster* stock. We would also like to thank Dr Todd Schlenke for providing the Lh14 and Lb17 wasp strains and Dr Alexandre Leitão for providing the NSRef wasp strain. We would also like to thank Dr Steve Parratt and Ben Walsh for collecting the wasp strain from Madeira. We also thank Prof Mariana Mateos and Paulino Ramirez for the *Spiroplasma* strain primer sequences. We thank Dr Ewa Chrostek, Prof Mariana Mateos and an anonymous reviewer for helpful comments on the manuscript. This project was supported by funding from the NERC (Studentship to JJ, grant number NE/L002450/1).

## Data accessibility

Data generated and analysed during this study are available at figshare (https://doi.org/10.6084/m9.figshare.c.4856790.v1).

## Supplementary material

**Table S1:**
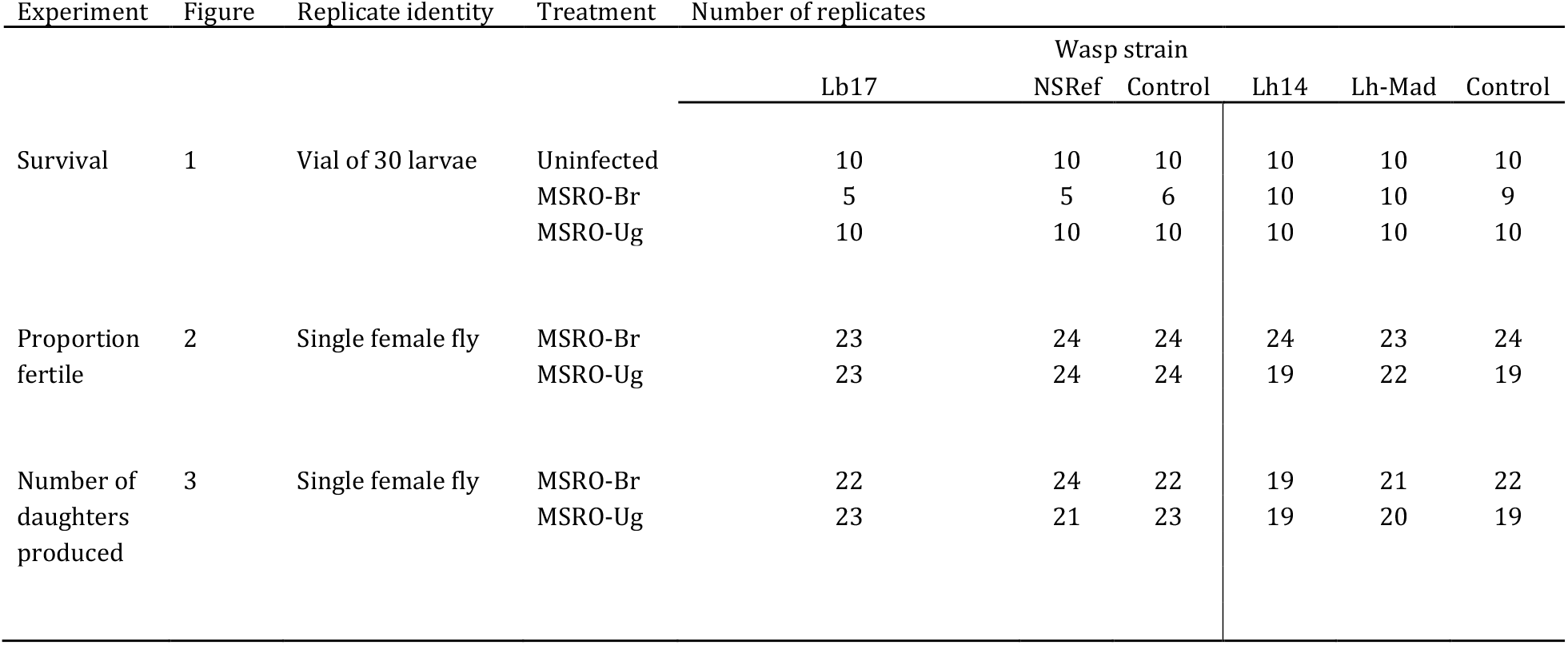
Replicate identity and number.

